# Introducing non-enzymatic crosslinks into atomistic simulations of collagen fibrils

**DOI:** 10.64898/2026.03.13.711566

**Authors:** Guido Giannetti, Justin Pils, Frauke Gräter, Debora Monego, Christoph Dellago

## Abstract

**Motivation:** Collagen fibrils are the primary load-bearing units of connective tissues. However, generating atomistic, simulation-ready models remains challenging due to collagen’s hierarchical organization and the diversity of its crosslinking network across tissues, ages, and metabolic states. Notably, non-enzymatic advanced glycation end-product (AGE) crosslinks—central to aging and diabetic complications—are largely absent from current atomistic fibril modelling workflows.

**Results:** Here, we present an extension of the ColBuilder framework to generate atomistic collagen fibril models that incorporate three representative AGE-derived crosslinks (glucosepane, pentosidine, and MOLD) alongside enzymatic crosslinks. Amber99-compatible parameters are provided and assessed against QM-optimized reference geometries using all-atom molecular dynamics (MD) simulations. As proof-of-concept, we examine the mechanical response of single D-period collagen microfibrils featuring enzymatic-only, AGE-only, and mixed crosslink patterns in Molecular Dynamics simulations under force, and observe that AGE crosslinks differently impact the fibril structure compared to enzymatic crosslinks. The extension to ColBuilder can aid future structure-based research on collagen aging.

**Availability and implementation:** ColBuilder is available as an open-source Python command-line package at https://github.com/graeter-group/colbuilder.

## 1 Introduction

Collagen is the principal structural protein of the extracellular matrix and provides the load-bearing capacity of connective tissues such as skin, bone, tendons, cartilage, and blood vessels. Collagen molecules assemble into D-periodic fibrils characterized by alternating overlap and gap regions, where the mechanical response is shaped by both molecular packing and the covalent crosslinks that interconnect neighbouring triple helices. Variations in crosslink chemistry, tissue specificity, and density modulate stiffness and viscoelasticity of collagen, thereby tuning physiological function and contributing to disease onset [1, 2]. Crosslinking is initiated enzymatically by lysyl oxidases (LOXs) through oxidative deamination of specific lysine and hydroxylysine residues, predominantly within the non-helical telopeptides. The resulting intermolecular covalent crosslinks bridge adjacent triple helices, reinforce the fibrillar network, and enhance tensile strength [2].

Owing to its exceptionally slow turnover, ranging from 1–2 years in bone to over 100 years in cartilage [3], collagen is also exposed to progressive, non-enzymatic glycation of lysine and arginine side chains through Maillard chemistry. In contrast to enzymatic crosslinking, which targets specific telopeptide–helix interfaces, glycation can occur along the collagen molecule and produces a chemically diverse spectrum of advanced glycation end-products (AGEs), including intramolecular adducts and intermolecular crosslinks. AGE accumulation alters fibril mechanics, typically increasing stiffness and promoting brittle failure. Moreover, AGEs can modify collagen hydration [4], a key determinant of collagen stability and self-assembly [5, 6], and impair molecular recognition by perturbing interactions with proteoglycans, cell-surface integrins, and collagenolytic enzymes [3, 7].

Despite major advances, high-resolution experimental structures remain largely restricted to collagen-mimetic peptides (CMPs), which have revealed the triple-helical conformation and key hydration features [8–10]. In contrast, native fibrils have proven much more difficult to characterize at atomic resolution [11–14], owing to their heterogeneity, flexibility, and dynamic packing [5, 15].

Atomistic modelling and molecular dynamics (MD) simulations provide a powerful complementary approach, enabling systematic exploration of fibril structure and crosslink chemistry [16–19]. However, current atomistic models largely focus on enzymatic modifications, leaving AGE-derived crosslink networks comparatively unexplored. To address this gap, we extended the open-source framework ColBuilder [20] to construct collagen fibrils that incorporate both intermolecular enzymatic crosslinks and three key AGE crosslink types: glucosepane, pentosidine, and MOLD (Figure 1). Glucosepane is widely reported as a major AGE crosslink in collagen and the extracellular matrix (ECM). Pentosidine is a long-established AGE crosslink that is routinely quantified and often used as a surrogate marker of glycation-related crosslinking in collagen-rich tissues. MOLD reflects the formation of methylglyoxal-driven cross-links associated with metabolic stress [2].

**Figure 1.**
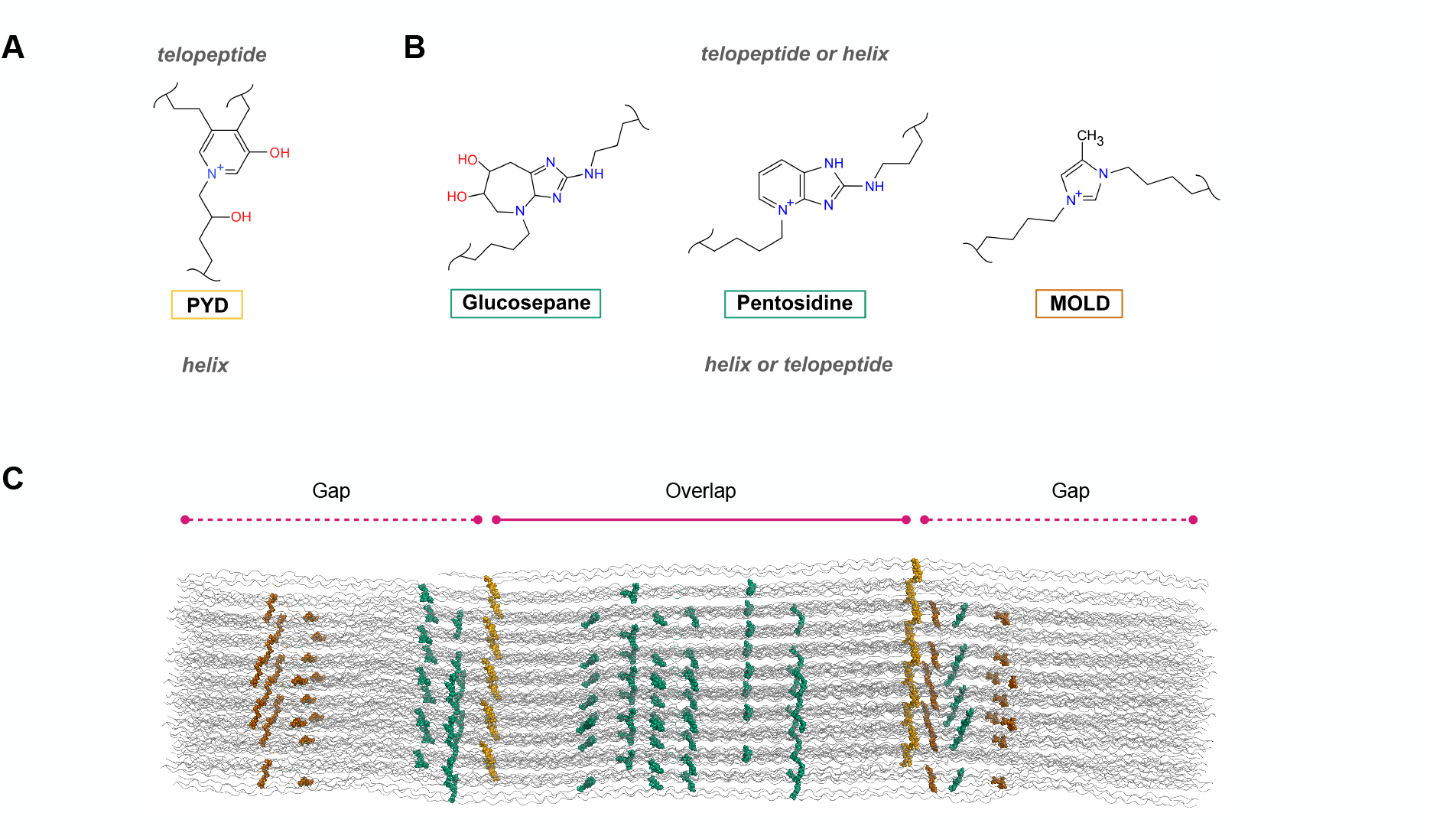
**A–B** Chemical structures of the crosslinks considered in this work. Pyridinoline (PYD), already implemented in [23], is a trivalent *enzymatic* crosslink that couples two residues in the telopeptide region of one collagen molecule to a residue in the triple-helical domain of a neighbouring molecule. In contrast, the AGE-derived crosslinks introduced here—glucosepane, pentosidine, and MOLD—are not confined to the canonical telopeptide–helix interface and can involve reactive residues distributed along the collagen sequence. **C** Schematic of a collagen D-period (∼67 nm) fibril highlighting crosslinking site diversity. Enzymatic PYD sites at the N- and C-terminal telopeptides are shown in amber. Candidate AGE sites are shown as Lys–Arg positions (glucosepane/pentosidine; green) and Lys–Lys positions (MOLD; rust). Sites shared by multiple crosslink types and duplicated across chains within the same triple helix are omitted for clarity; the schematic is intended to emphasize the broader set of admissible AGE crosslink locations relative to enzymatic crosslinks.

The extended framework allows for systematic control over fibril composition and geometry. Users can specify crosslink combinations and density, as well as fibril diameter and length across 19 vertebrate collagen sequences. To enable downstream simulations, we provide Amber99-compatible parameters for the three AGE crosslinks, and we use all-atom MD on a single D-period fibril with distinct crosslinking networks to illustrate their stability and mechanical impact.

The rise in life expectancy and widespread hyperglycemia in type II diabetes drive the accumulation of AGE, making glycation-induced changes in the extracellular matrix a growing health concern. In this context, our atomistic AGE-inclusive collagen fibrils provide a versatile platform for investigating the molecular consequences of glycation through multiscale simulations. They also provide high-resolution templates to support cryo-EM fitting, enable structure-based design efforts, and ultimately deepen our understanding of how aging and metabolic stress remodel collagen’s mechanical properties and biological interactions.

## 2 Materials and methods

ColBuilder [20] generates simulation-ready type I collagen microfibril models by combining triple-helix structure generation through homology modelling, crystallography-informed fibril assembly, and automated topology generation for enzymatic crosslinks. Here we extend this framework to incorporate AGE-derived crosslinks through three key developments: (1) expanding the crosslink database to include three AGE crosslinks (glucosepane, MOLD, and pentosidine) and their respective candidate sites in collagen type I fibrils, (2) implementing sequential crosslink insertion with flexible periodic neighbour selection during crosslink optimization, and (3) developing Amber99-compatible parameters for the AGEs added.

### 2.1 Candidate AGE crosslink positions

Enzymatic collagen crosslinks arise at well-defined intermolecular interfaces, coupling reactive residues in the N- and C-terminal telopeptides to residues in the triple-helical domain of an adjacent molecule. By contrast, AGE-derived crosslinks can involve residues distributed throughout the collagen sequence, thereby expanding the set of admissible crosslinking sites within the fibril. The chemistries considered here and the broader distribution of candidate AGE sites relative to PYD are summarized in Figure 1.

To derive candidate AGE crosslink positions, we analyzed 19 vertebrate collagen type I sequences spanning mammals, fish, and reptiles (Supplementary Table S2). For each sequence, we generated a periodic fibril representation by expanding a full-length collagen triple helix of approximately 300 nm into symmetry-related neighbours of the crystallographic unit cell. This was performed by applying crystal symmetry operations and unit-cell translations within a user-defined contact-distance cutoff using UCSF Chimera [21]. We then pre-screened candidate residue pairs with an in-house MDAnalysis [22] workflow by applying a distance-based geometric filter, consistent with prior *in silico* approaches for identifying candidate glycation-derived crosslink sites [7]. Specifically, we enumerated all Lys–Arg (for glucosepane and pentosidine) and Lys–Lys (for MOLD) pairs and retained those with C*α*–C*α* distances at or below 15 Å. We chose this conservative cutoff to avoid excluding plausible sites given the limited experimental data on AGE crosslink locations in native fibrils. For example, in *Rattus norvegicus* (the template sequence), this geometric screening identified 102 Lys–Arg candidate sites (for glucosepane/pentosidine) and 45 Lys–Lys candidate sites (for MOLD). Across all 19 species, candidates ranged from 77 to 107 sites for glucosepane/pentosidine and from 36 to 45 for MOLD. The retained candidates were subsequently evaluated by explicit model building, with successful refinement achieved for approximately 64% of the candidates as described below.

### 2.2 AGE insertion and refinement

To support AGE-derived crosslinks, we extended ColBuilder’s sequence-generation module to enable generation of purely AGE-crosslinked fibrils or fibrils combining both enzymatic and AGE crosslinks. For combined crosslinking, a staged procedure is required. From a FASTA input, ColBuilder first generates an enzymatically crosslinked triple helix following the workflow described previously [20]. This enzymatically crosslinked structure then serves as input for a second pass, in which AGE crosslink markers are introduced via MODELLER patching [24] and refined by optimizing only the newly added AGE constraints. This sequential approach prevents interference between crosslink types during optimization and supports iterative extension, allowing additional AGE-derived crosslinks to be introduced in further cycles without repeating the initial enzymatic crosslinking step.

A central methodological change, as illustrated in Figure 2, addresses the distinct spatial requirements of enzymatic versus AGE crosslinks. Enzymatic crosslinks form at specific telopeptide-to-helix interfaces, allowing the original ColBuilder to use a single fixed periodic neighbour for all crosslink refinement and optimization. In contrast, AGE crosslinks can form between residues located anywhere along the triple helix. Because the collagen fibril has a staggered D-periodic structure, residues at different axial positions have different intermolecular neighbours; a Lys-Arg pair in the gap region requires a different periodic neighbour than a pair in the overlap region, for example. We therefore implemented user-selectable unit-cell translations, enabling each AGE crosslink site to be refined against its geometrically appropriate neighbour. During refinement, the fibrillar context is approximated by a minimal periodic environment comprising the target triple helix plus one periodic image selected by the specified unit-cell translation vector, and crosslink-forming atom distances are evaluated across this molecular pair.

**Figure 2.**
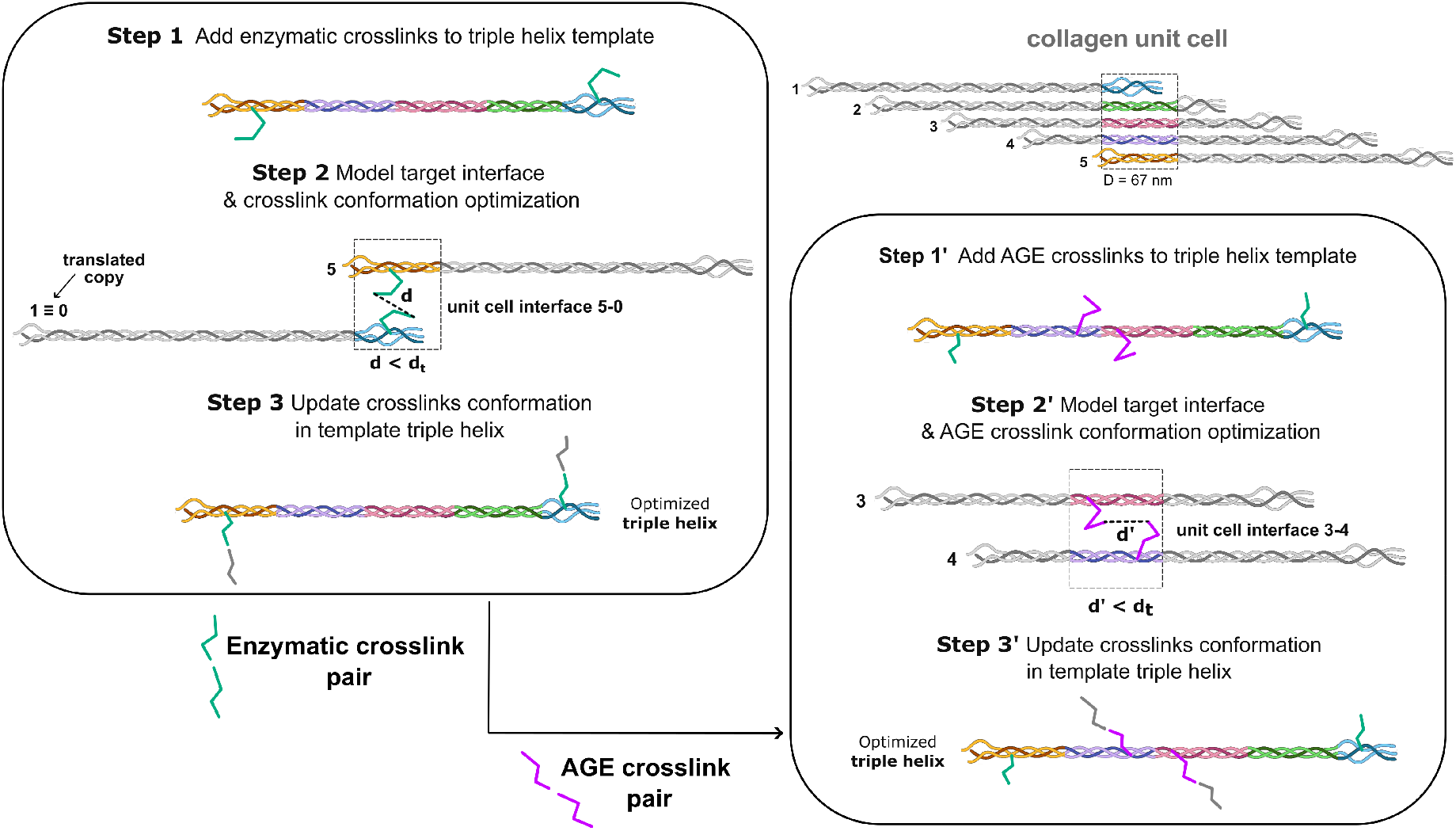
Staged, interface-specific refinement of enzymatic and AGE-derived crosslinks in ColBuilder. **Left panel (enzymatic refinement):** Enzymatic crosslinks are refined by modelling a single, predefined telopeptide– helix interface (Step 2). **Right panel (AGE extension):** Starting from an enzymatically crosslinked template, AGE crosslinks—which can occur at different axial positions along the collagen molecule—are introduced in a second pass (Step 1^*′*^) and refined against a site-specific periodic neighbour (illustrated by distinct unit-cell interfaces; Step 2^*′*^). Candidate sites are accepted if the refined geometry satisfies a distance criterion (*d < d*_*t*_, here *d*_*t*_ = 5 Å) and is free of steric clashes. The optimized crosslink geometry is propagated back onto the template triple helix (Steps 3/3^*′*^). The unit-cell schematic **(top)** highlights the staggered molecular packing over one D-period (∼67 nm), motivating the need for different interfaces depending on crosslink location.

All geometrically pre-screened candidates were carried forward to explicit model building. For each candidate site, we built the AGE structure via MODELLER patching and performed local refinement using the site-specific periodic neighbour. We evaluated refinement success based on absence of steric clashes and maximum crosslink-forming atom separation of 5 Å(we note that these were further optimized in a subsequent energy minimization step for realistic bond lengths). Successfully refined sites were stored in the extended crosslink database, where we recorded the AGE chemistry type, participating atom identities, and corresponding unit-cell translation vectors, enabling users to generate highly customizable collagen microfibrils incorporating both enzymatic and AGE-derived crosslinks.

### 2.3 Parameterization of AGE crosslinks

Amber99-compatible force-field parameters for the three AGE-derived crosslinks were generated as follows. First, the crosslink structures were built and capped with acetyl and N-methyl groups using Maestro [25]. Second, geometries were optimized and the electrostatic potential was computed at the B3LYP/6-31G* level in Gaussian [26–28]. Third, partial atomic charges were derived by restrained electrostatic potential fitting using Antechamber [29, 30]. During fitting, atoms belonging to the capping groups were constrained to reproduce the corresponding AMBER99 charges [31]. The resulting parameters were converted to GROMACS format using Acpype [32].

To minimize the introduction of new parameters and maintain compatibility with the existing force field, we retained only those required to describe the new covalent connectivity introduced by the AGE-derived crosslinks and not already defined in Amber99SB*-ildnp [33]. All newly introduced parameters, such as cross-link topologies, special bonds, partial charges, and bonded parameters, provided in Supplementary Informations, Figures S2–S4 and Tables S3–S9.

### 2.4 Crosslink specification for enzymatic- and AGE-crosslinked microfibrils

ColBuilder enables user-defined control over crosslink chemistry, mixing, and density while preserving a crystallography-informed fibrillar architecture. The current implementation extends the original functionality by supporting AGE-derived crosslinks and user-prescribed mixtures of enzymatic and non-enzymatic crosslink types.

Two complementary strategies are provided. First, heterogeneous microfibrils can be generated by mixing pre-built triple-helix variants that carry different crosslink sets at user-defined ratios, thereby controlling the fraction of distinct enzymatic and AGE-bearing populations within the same assembly. Second, crosslink density can be reduced by replacing a user-defined fraction of crosslink markers with their native amino acids using UCSF Chimera’s `swapaa` command [21]. This replacement can be applied selectively to enzymatic markers, AGE markers, or both, and restores the corresponding native residue identity (Lys or Arg), depending on the crosslink chemistry.

These capabilities support controlled model sets in which crosslink chemistry and density are varied systematically, facilitating direct in silico comparisons of their effects on fibril structure and mechanics. We applied these strategies to generate five distinct crosslink configurations for validation: pure PYD (enzymatic control), pure glucosepane, pure MOLD, pure pentosidine, and mixed glucosepane+PYD. In each case, we chose the maximal crosslink density for the fibril at hand. Detailed usage examples are provided in the ColBuilder GitHub repository.

## 3 Results

### 3.1 Assessment of AGE crosslink parameters

Although the AGE-derived crosslinks were parameterized using an established RESP-based workflow, experimental reference data for a direct validation of these parameters are not available to our knowledge. We therefore benchmarked the Antechamber/RESP parametrization against machine-learned parameters generated with Grappa [34]. For each AGE crosslink and parameter set, we ran five independent 100-ns replicas (3.0 µs total sampling across the three crosslinks and both parameter sets) using GROMACS 2024.4 [35]. We assessed agreement with the QM reference geometry used for RESP fitting by comparing ring-core angle distributions (i.e., excluding junction angles connecting the ring system to side-chain atoms and ring substituents, highlighted in yellow in Table S10) and complementary structure-based metrics (RMSD, RMSF, and end-to-end distance; Figure S5).

Across all ring-core angles (27 angles over the three AGEs), Antechamber reproduces the QM reference with a mean absolute deviation of 2.72^*°*^, with 85% of angles within 5^*°*^ of the QM values. Grappa shows a closer match, with a mean absolute deviation of 1.05^*°*^, and all ring-core angles within 5^*°*^ of the QM reference. The MD angle distributions display similar widths for the two parameterizations, with average standard deviations of approximately 2.5^*°*^. Representative snapshots comparing parameter sets are shown in Figure 3.

**Figure 3.**
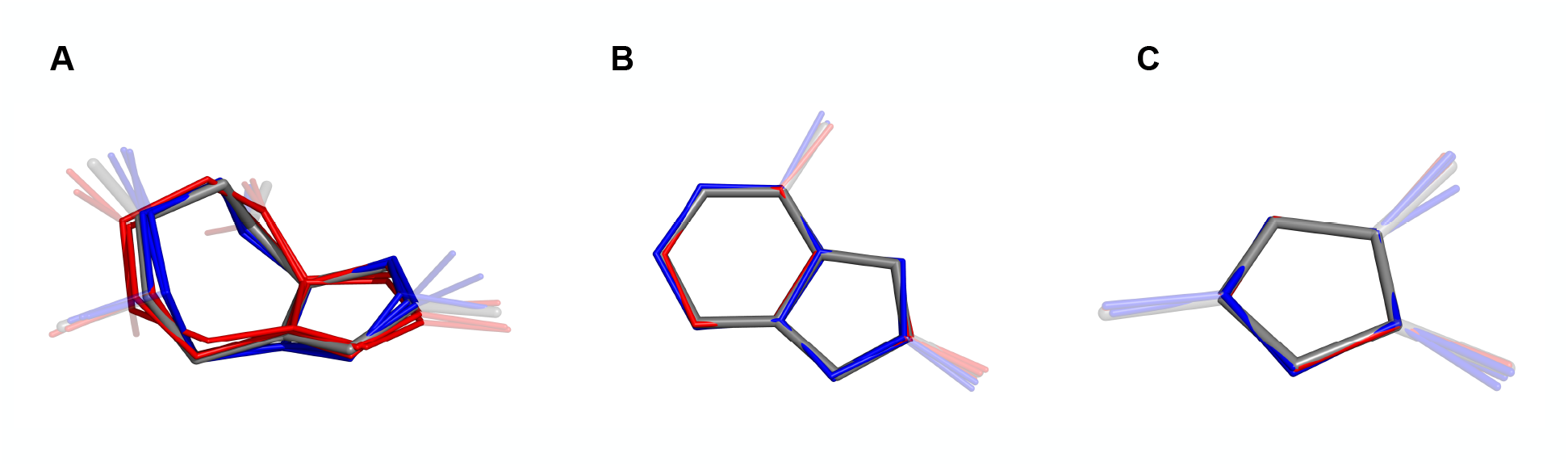
Representative MD snapshots of the three AGE-derived crosslinks: **A** glucosepane, **B** pentosidine, and **C** MOLD. The QM reference geometry used for RESP fitting is shown in grey, Grappa-derived parameters in red, and Antechamber/RESP-derived parameters in blue. Snapshots were taken at 50, 75, and 100 ns from a single arbitrarily selected replica (visualization only). Opaque bonds highlight the ring-core atoms used for ring-only angle, RMSD, and RMSF analyses, whereas translucent bonds indicate connections to side-chain atoms and ring substituents (C–OH and C–CH_3_ substituents in glucosepane and MOLD, respectively. Hydrogen atoms are omitted for clarity).

Angle-level agreement is further supported by structural metrics referenced to the QM geometry. All-atom RMSD values are nearly identical for the Antechamber and Grappa parameterizations across all three crosslinks (Antechamber: 0.491–0.533 nm; Grappa: 0.490–0.537 nm). Restricting the RMSD to ring atoms reduces the values by about four-fold, to 0.124–0.139 nm (Antechamber) and 0.126–0.135 nm (Grappa), indicating that the ring-core geometry remains tightly preserved under both parameterizations and that the side-chain moieties dominate the overall deviation. RMSF follows the same trend: mean all-atom RMSF values are comparable between parameter sets (Antechamber: 0.287–0.310 nm; Grappa: 0.297–0.317 nm), while ring-atom RMSF values are lower (Antechamber: 0.254–0.277 nm; Grappa: 0.242–0.275 nm). Finally, the end-to-end distance—defined between the terminal C*α* atoms of each crosslink and therefore sensitive to side-chain conformational sampling—exhibits the strongest parameter dependence, most prominently for pentosidine and MOLD, for which Grappa yields smaller mean values than Antechamber.

Taken together, these results show that, although Grappa yields a slightly closer match to the QM reference for ring-core angular geometry—as expected for a machine-learned parametrization framework trained directly on QM reference data to reproduce energies and forces—the RESP/Antechamber-derived parameters provide a robust description of the AGE ring-core geometry and remain stable over the simulated timescales. This supports their use as a consistent extension of the established ColBuilder parameterization strategy previously adopted for enzymatic crosslinks [23].

### 3.2 Collagen microfibril all-atom simulations

To assess the structural integrity and tensile response of ColBuilder-generated models, we performed all-atom molecular dynamics simulations under constant-force loading. To specifically probe the effect of AGE-derived crosslinks on fibril mechanics, we simulated a single D-period (∼67 nm), spanning one overlap and one gap region, with different crosslinking patterns.

Microfibril topologies were generated automatically within the ColBuilder workflow using the GROMACS [35] tool `pdb2gmx`, with the Amber99SB*-ildnp force field complemented with the AGE parameters described above [33, 36]. The resulting solvated systems consisted of approximately 3.5 × 10^6^ atoms using the TIP3P water model. Simulations were run with GROMACS 2024.4/2024.2 on MUSICA (Austrian Scientific Cluster) and LEONARDO (CINECA), achieving ∼ 12 ns/day.

Each system was first minimized in vacuum, then solvated and minimized again. Equilibration was performed in two stages with gradually increased time steps from 0.5 to 2 fs: (i) 10 ns in the NVT ensemble, followed by (ii) 5 ns in the NPT ensemble. Temperature and pressure were maintained at 300 K and 1 atm, respectively, using the stochastic velocity–rescaling thermostat [37] and the Parrinello–Rahman barostat [38]. To preserve the fibrillar arrangement while enabling local relaxation, heavy atoms were position-restrained throughout equilibration; position-restraint force constants were decreased stepwise from 2500 to 1000 kJ mol^−1^ nm^−2^ during NVT and from 1000 to 300 kJ mol^−1^ nm^−2^ during NPT. Periodic boundary conditions were applied in all directions.

The loading force was applied along the fibril axis between the centers of mass of the terminal ACE and NME cap methyl groups and increased stepwise to allow structural adaptation. The load was ramped from 0 to 1 nN per strand in 0.2 nN increments over 12.5 ns and then held for 87.5 ns (100 ns total). This target force was chosen to be consistent with estimated physiological loads on single collagen molecules (corresponding to stresses in the tens-of-MPa range) [39], while the simulation length was selected to capture the early stages of force-induced reorganization reported in previous atomistic studies [16].

To prevent force-induced unwinding of the triple helices, all simulations included torque restraints implemented with the GROMACS enforced-rotation framework. Rotation groups were defined for each strand using the methyl atoms of the ACE and NME terminal caps, and the corresponding restraint blocks were generated automatically with an in-house script. The restraints acted about the fibril axis, enforced zero net twist, and used a harmonic potential with a force constant of 200 kJ mol^−1^ nm^−2^.

We considered the following crosslink setups: (i) a reference system containing only enzymatic PYD crosslinks at both the N- and C-terminal telopeptide sites; (ii) three AGE-only variants in which the terminal enzymatic crosslinks were replaced by one of the three AGE chemistries (glucosepane, pentosidine, or MOLD), implemented at both termini in each system; and (iii) a mixed system combining terminal PYD crosslinks with an additional glucosepane crosslink within the overlap region (Figure 4C).

**Figure 4.**
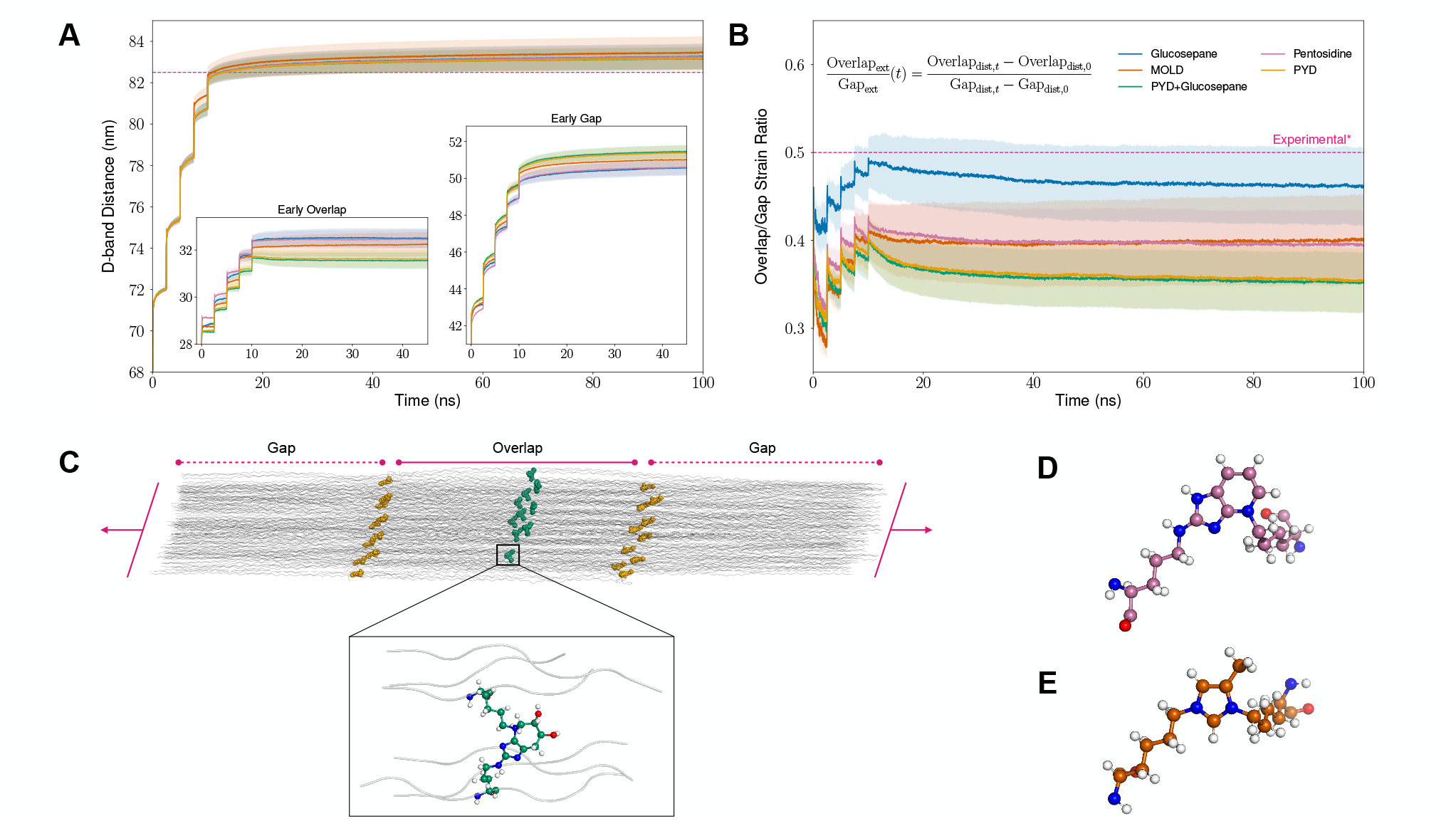
Validation of differently crosslinked collagen microfibrils by all-atom constant-force MD simulations. **(A)** Time evolution of the end-to-end distance of a single-D-period microfibril (D-band proxy), with insets showing the corresponding overlap and gap contributions at early simulation times. **(B)** Time evolution of the overlap-to-gap strain ratio. In both panels, magenta dashed lines mark experimentally reported reference values [40], while shaded regions indicate the standard error of the mean of replica-averaged observables, estimated by combining frame-wise variability with trajectory-scale uncertainty (see Supplementary Methods). **(C)** Representative snapshot of the mixed PYD+glucosepane system under load, with terminal PYD crosslinks (amber spheres) and an additional glucosepane crosslink in the overlap region (green spheres; inset). No individual collagen molecule is pulled out during the simulation, indicating that microfibril integrity is preserved under the applied force. Distinct deformation modes are evident: the overlap region adopts a more parallel arrangement of neighbouring triple helices and exhibits less elongation, whereas the gap region remains more intertwined and undergoes greater stretching. **(D–E)** Molecular structures of pentosidine and MOLD crosslinks, respectively (ball-and-stick; carbon colouring as in **(A–B)**); capping groups are omitted for clarity.

Mechanical response was characterized at the fibril scale by monitoring the end-to-end distance of the simulated D-period, used here as a proxy for D-band extension in this reduced system, together with the extensions of the overlap and gap regions and the overlap-to-gap strain ratio. To ensure reproducibility and assess variability, each crosslink configuration was simulated in three independent replicas.

All fibril-level observables displayed a two-stage response, with a rapid elongation during force ramping followed by an asymptotic approach to a plateau value (Figure 4A). The single D-period, defined as the sum of the overlap and gap regions, elongated by approximately 24% overall. Final extensions fell within a narrow range across all crosslinking patterns, from 83.15 nm for the mixed PYD+glucosepane and PYD-only systems to 83.45 nm for the AGE-only MOLD configuration, with the corresponding AGE-only glucosepane and pentosidine systems adopting intermediate values. Larger differences emerged in the relative contributions of the overlap and gap regions, which varied in opposite directions across the different crosslinking patterns (Figure 4A, insets). Across all crosslinking patterns, the overlap increased only marginally, from about 27.4–27.9 nm to roughly 31.6–32.5 nm, whereas the gap extended from about 39.2–39.8 nm to 50.7–51.6 nm. Our results are consistent with previous all-atom constant-force simulations and AFM nanoindentation experiments, which reported D-period extensions of 82.5 *±* 1.0 nm and 80–82.5 nm, respectively, as well as a 25–100% higher stiffness in the overlap region compared to the gap region [17, 20, 40–42]. Within the observed extensions of overlap and gap regions, the AGE-only systems generally showed comparably large extensions for the overlap region, whereas the PYD-only reference and the mixed PYD+glucosepane system extended less within their overlap regions; the opposite trend was observed for the gap region.

Adding glucosepane crosslinks within the overlap region did not measurably alter the mechanical response relative to the PYD-only reference system. By contrast, replacing the native terminal PYD crosslinks with AGE-derived ones produced a modest but systematic redistribution of deformation, shifting extension toward the overlap region and away from the gap region without appreciably altering the overall D-period elongation. These findings indicate that AGE effects emerge more clearly from replacement of key load-bearing enzymatic crosslinks than from simply increasing crosslink density. Our results also underline that enzymatic and non-enzymatic crosslinks, with their different chemical structure and their specific localization within the fibril topology, effect the internal force distribution in loaded collagen fibrils very distinctly. How AGE crosslinks impact collagen rupture and mechanoradical formation [17] remains to be explored.

Overall, the constant-force simulations indicate that ColBuilder-generated models preserve structural integrity under load across all crosslink chemistries. This robustness extends beyond enzymatically crosslinked microfibrils (PYD) to AGE-only architectures (glucosepane, pentosidine, MOLD) and to mixed enzymatic/non-enzymatic networks, supporting the use of the extended ColBuilder framework for mechanistic studies of how crosslink chemistry modulates collagen mechanics at the fibrillar scale.

## 4 Conclusions

In this work, we extended the ColBuilder framework to enable atomistic modelling of collagen fibrils that combine enzymatic crosslinks with representative AGE-derived chemistries. The implementation expands the crosslink database and introduces a staged insertion strategy that supports iterative refinement of additional crosslinks on pre-crosslinked scaffolds, together with user control over the periodic neighbour used during crosslink optimization. To make these models directly usable in molecular simulations, we developed Amber99-compatible parameters for glucosepane, pentosidine, and MOLD and integrated them into the ColBuilder topology-generation pipeline.

Benchmark simulations of the isolated AGE motifs indicate that the Antechamber-derived parameters preserve the ring-core geometry and provide a consistent extension of the enzymatic-crosslink parameterization strategy. In constant-force simulations of a single D-period collagen microfibril, AGE-inclusive models remain mechanically stable and reproduce the characteristic differential deformation of the overlap and gap regions. AGE-related effects at the fibrillar scale emerge more clearly when non-enzymatic crosslinks replace key load-bearing enzymatic crosslinks than when they simply increase crosslink density.

Beyond the specific crosslink patterns examined here, ColBuilder supports user-defined control over crosslink density and mixed enzymatic/AGE networks, enabling systematic *in silico* studies of age- and tissue-dependent remodeling as enzymatic crosslinking varies and AGEs accumulate. We also provide guidelines to straightforwardly extend ColBuilder to other collagen crosslinks or post-translational modifications of interest. As new experimental and computational inputs on collagen hierarchical structure become available—including electron microscopy reconstructions, updated crosslinking data (particularly for AGE-derived and other post-translational modifications) from mass spectrometry, and complementary structural insights from prediction approaches such as AlphaFold [43]— ColBuilder provides a natural route to incorporate these data into simulation-ready fibril models.

## Competing interests

No competing interest is declared

## Author contributions statement

Guido Giannetti (Conceptualization [lead], Software [equal], Data curation [lead], Methodology [equal], Formal analysis [lead], Investigation [lead], Validation [lead], Writing—original draft [lead], Writing—review & editing [equal]), Debora Monego (Conceptualization [supporting], Software [equal], Data curation [supporting], Methodology [equal], Formal analysis [supporting], Investigation [supporting], Validation [supporting], Writing—original draft [supporting], Writing—review & editing [equal]), Justin Pils (Conceptualization [supporting], Software [supporting], Data curation [supporting], Methodology [supporting], Formal analysis [supporting], Investigation [supporting], Validation [supporting], Writing—original draft [supporting], Writing—review & editing [supporting]), Frauke Gräter (Conceptualization [supporting], Methodology [supporting], Writing—original draft [supporting], Writing—review & editing [equal]), and Christoph Dellago (Conceptualization [supporting], Methodology [supporting], Funding acquisition, Writing—original draft [supporting], Writing—review & editing [equal])

## Data Availability Statement

FASTA files for collagen type I sequences used in this study are available in the ColBuilder GitHub repository. The repository at https://github.com/gggguido/AGEs-ColBuilder.git provides the simulation input files used for the fibril simulations, including GROMACS . `mdp` files for equilibration and production runs; scripts for trajectory and structural analyses; representative equilibrated fibril structures for the AGE-only, enzymatic-only, and mixed systems; scripts for identifying candidate crosslinking sites in the fibril models; and a utility for generating MODELLER internal-coordinate (IC) lists from crosslink SMILES strings. The repository further includes the topology files used for the isolated AGE-crosslink simulations in water, specifically the `topol.top` files based on Antechamber-derived parameters and the `grappa topol.top` files containing Grappa-derived parameters, together with a guide and example files for parameterizing additional Amber99-compatible crosslinks.

## Acknowledgments

G. G. acknowledges financial support from the Austrian Science Fund (FWF) under project P35715 and from the Vienna Doctoral School in Physics (VDSP).

Computational results were obtained using the Austrian Scientific Computing (ASC) infrastructure and the LEONARDO supercomputer at CINECA (Italy) via AURELEO (Austrian Users at LEONARDO) projects n. 72451 and n. 72829.

## List of Supplementary Materials

All data are incorporated into the article and its online supplementary material.

## Notes

https://github.com/gggguido/AGEs-ColBuilder.git

https://github.com/graeter-group/colbuilder

### Competing Interest Statement

The authors have declared no competing interest.

