## Supplementary Information for "Introducing non-enzymatic crosslinks into atomistic simulations of collagen fibrils"

#### 1 Crosslinks

ColBuilder encodes each intermolecular crosslink in a unified format by specifying the modified residues involved and the corresponding atom–atom connection points. These connection points are highlighted in cyan in this document and are reported as *special bonds* using the standard GROMACS nomenclature. Enzymatic crosslinks typically involve Lys-derived marker residues and are represented by two (divalent) or three (trivalent) connection points. AGE-derived crosslinks involve either Lys–Arg or Lys–Lys residue pairs, are represented by two connection points, and can be specified at sites distributed along the collagen sequence rather than being restricted to the N- and C-terminal regions.

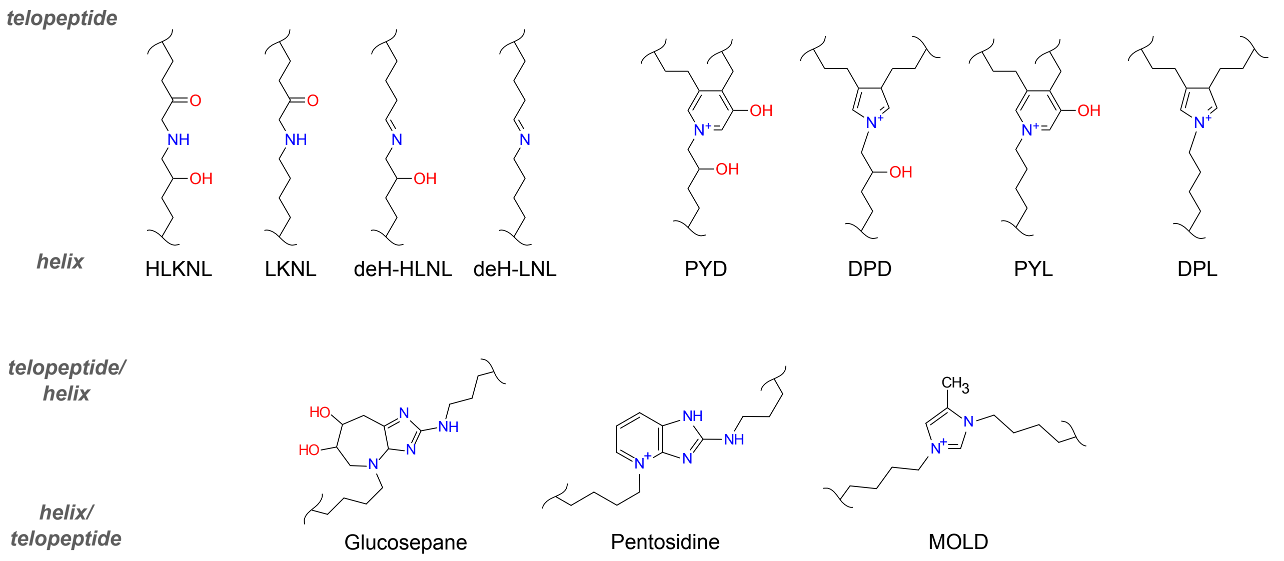


**Figure S1.** The first row shows the enzymatic crosslink already implemented in [1], which forms at well-defined intermolecular interfaces by coupling a residue in the telopeptide of one collagen molecule to a residue in the triple-helical domain of an adjacent molecule (schematized as telopeptide/helix). The second row shows the three non-enzymatic AGE crosslinks introduced in this work—glucosepane and pentosidine (Lys–Arg-derived) and MOLD (Lys–Lys-derived). These are not restricted to specific collagen regions (schematized as telopeptide/helix or helix/telopeptide), reflecting the fact that AGE formation can in principle involve residues distributed along the full collagen sequence.

#### 2 Crosslinks database (`crosslinks.csv`)

| OR | term | combination | type | R1 | A1 | P1 | R2 | A2 | P2 | R3 | A31 | A32 | P3 | #*shift* |
|---|---|---|---|---|---|---|---|---|---|---|---|---|---|---|
| ARG_LYS | N | 310.C – 542.B | Glucosepane | AGS | NZ | 310.C | LGX | CE | 542.B | NONE | NONE | NONE | NONE | #D3–D4 |
| ARG_LYS | N | 509.B - 280.A | Pentosidine | APD | NZ | 509.B | LPS | CE | 280.A | NONE | NONE | NONE | NONE | #D2–D3 |
| LYS | N | 597.A - 358.A | MOLD | LZS | NZ1 | 597.A | LZD | CE | 358.A | NONE | NONE | NONE | NONE | #D3–D4 |

**Table S1.** Example of the three AGE-derived crosslink entries in ColBuilder for *Rattus norvegicus*, the template sequence. Columns report the original residue pair (OR), the terminal label (term; required by ColBuilder even for non-terminal AGE sites), the residue pair used for placement (combination), the crosslink chemistry (type), and the residue/atom/position triplets defining the reactive connection points (R1/A1/P1 and R2/A2/P2). The third triplet (R3/A31/A32/P3) is used only for trivalent crosslinks and is set to `NONE` for divalent and AGE-derived crosslinks. Residue positions are given as *residue.chain*. The last column records the required unit cell translation.

#### 3 Number of AGE-derived crosslinks

| Species | # Lys_Arg total | # Lys_Arg retained | # Lys_Lys total | # Lys_Lys retained |
|---|---|---|---|---|
| *ailuropoda melanoleuca* | 88 | 58 | 43 | 24 |
| *bos taurus* | 104 | 64 | 41 | 24 |
| *callithrix jacchus* | 98 | 63 | 39 | 20 |
| *canis lupus* | 104 | 65 | 40 | 20 |
| *danio rerio* | 107 | 63 | 36 | 18 |
| *homo sapiens* | 101 | 66 | 39 | 20 |
| *loxodonta africana* | 95 | 60 | 43 | 21 |
| *mus musculus* | 101 | 61 | 42 | 25 |
| *mustela putorius* | 101 | 67 | 39 | 20 |
| *myotis lucifugus* | 105 | 67 | 41 | 23 |
| *oreochromis niloticus* | 106 | 65 | 36 | 19 |
| *oryzias latipes* | 106 | 65 | 36 | 19 |
| *otolemur garnettii* | 102 | 65 | 41 | 24 |
| *pan troglodytes* | 103 | 65 | 41 | 25 |
| *pelodiscus sinensis* | 97 | 66 | 40 | 16 |
| *pongo abelii* | 98 | 63 | 42 | 23 |
| *rattus norvegicus* | 102 | 64 | 45 | 27 |
| *tetraodon nigroviridis* | 77 | 47 | 42 | 27 |
| *xiphophorus maculatus* | 96 | 54 | 41 | 26 |

**Table S2.** Counts of candidate and retained AGE-derived crosslink sites identified across 19 vertebrate collagen type I sequences. Lys–Arg pairs correspond to candidate sites for glucosepane and pentosidine, whereas Lys–Lys pairs correspond to candidate sites for MOLD. "Total" reports the number of residue pairs passing the geometric pre-screening (C$\alpha$–C$\alpha$ $\leq$ 15 Å) in the periodic fibril representation, and "retained" reports the subset of candidates that, after explicit model building and local refinement within the fibril environment, satisfy the distance criterion (5 Å) between the reactive atoms forming the crosslink. All species names available in ColBuilder are listed in the first column, and the corresponding crosslink counts (total candidates and retained sites) are reported for each species in the remaining columns.

### 4 Residue topology and atom charges (*aminoacids.rtp*)

#### 4.1 Glucosepane crosslink

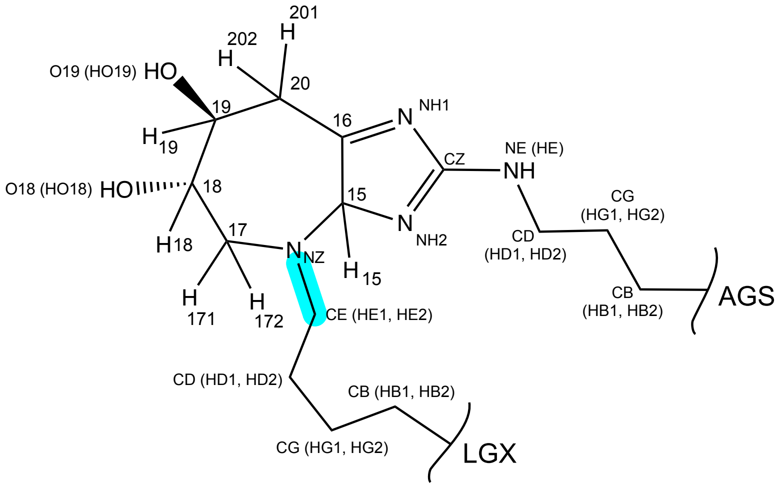


**Figure S2. Glucosepane crosslink.** Atom names and types as defined in the *aminoacids.rtp* file (in *GROMACS* format). The excerpt below shows the entries corresponding to Glucosepane (`LGX`: modified Lys residue & `AGS`: modified Arg residue).

[LGX]

[ atoms ]

| | | | |
|---|---|---|---|
| N | N | -0.448402 | 1 |
| H | H | 0.244307 | 2 |
| CA | CT | 0.219872 | 3 |
| HA | H1 | 0.060908 | 4 |
| CB | CT | -0.517274 | 5 |
| HB1 | HC | 0.160769 | 6 |
| HB2 | HC | 0.160769 | 7 |
| CG | CT | -0.046119 | 8 |
| HG1 | HC | 0.028325 | 9 |
| HG2 | HC | 0.028325 | 10 |
| CD | CT | 0.069291 | 11 |
| HD1 | HC | 0.031814 | 12 |
| HD2 | HC | 0.031814 | 13 |
| CE | CT | -0.453323 | 14 |
| HE1 | H1 | 0.168580 | 15 |
| HE2 | H1 | 0.168580 | 16 |
| C | C | 0.587484 | 18 |
| O | O | -0.561561 | 19 |

[ bonds ]

| | |
|---|---|
| N | H |
| N | CA |
| CA | HA |
| CA | CB |
| CA | C |
| CB | HB1 |
| CB | HB2 |
| CB | CG |
| CG | HG1 |
| CG | HG2 |
| CG | CD |
| CD | HD1 |
| CD | HD2 |
| CD | CE |
| CE | HE1 |
| CE | HE2 |
| C | O |
| -C | N |

[ impropers ]

| | | | |
|---|---|---|---|
| -C | CA | N | H |
| CA | +N | C | O |

#### [AGS]

[ atoms ]

| | | | |
|---|---|---|---|
| N | N | -0.409776 | 1 |
| H | H | 0.227143 | 2 |
| CA | CT | 0.134081 | 3 |
| HA | H1 | 0.036462 | 4 |
| CB | CT | -0.030064 | 5 |
| HB1 | HC | 0.026185 | 6 |
| HB2 | HC | 0.026185 | 7 |
| CG | CT | 0.024089 | 8 |
| HG1 | HC | -0.005728 | 9 |
| HG2 | HC | -0.005728 | 10 |
| CD | CT | 0.122941 | 11 |
| HD1 | H1 | 0.040543 | 12 |
| HD2 | H1 | 0.040543 | 13 |
| NE | N2 | -0.649385 | 14 |
| HE | H | 0.368975 | 15 |
| CZ | C | 0.791168 | 16 |
| NH1 | N* | -0.471221 | 17 |
| NH2 | N* | -0.897004 | 18 |
| C | C | 0.506324 | 19 |
| O | O | -0.523842 | 20 |
| C15 | CT | 0.707421 | 20 |
| H15 | H2 | -0.022832 | 21 |
| C16 | C | 0.019125 | 22 |
| NZ | N3 | -0.085682 | 17 |
| C17 | CT | -0.273185 | 23 |
| H171 | H1 | 0.111278 | 24 |
| H172 | H1 | 0.111278 | 25 |
| C18 | CT | 0.212995 | 26 |
| H18 | H1 | 0.106158 | 27 |
| O18 | OH | -0.643348 | 28 |
| HO18 | HO | 0.395352 | 29 |
| C19 | CT | 0.207711 | 30 |
| H19 | H1 | 0.003185 | 31 |
| O19 | OH | -0.614835 | 32 |
| HO19 | HO | 0.388586 | 33 |
| C20 | CT | -0.069824 | 34 |
| H201 | HC | 0.080283 | 35 |
| H202 | HC | 0.080283 | 36 |

[ bonds ]

| | |
|---|---|
| N | H |
| N | CA |
| CA | HA |
| CA | CB |
| CA | C |
| CB | HB1 |
| CB | HB2 |
| CB | CG |
| CG | HG1 |
| CG | HG2 |
| CG | CD |
| CD | HD1 |
| CD | HD2 |
| CD | NE |
| NE | HE |
| NE | CZ |
| CZ | NH1 |
| CZ | NH2 |
| NH2 | C15 |
| NH1 | C16 |
| C15 | H15 |
| C15 | NZ |
| C17 | NZ |
| C17 | H171 |
| C17 | H172 |
| C17 | C18 |
| C18 | H18 |
| C18 | O18 |
| O18 | HO18 |
| C18 | C19 |
| C19 | H19 |
| C19 | O19 |
| O19 | HO19 |
| C20 | C19 |
| C20 | H201 |
| C20 | H202 |
| C20 | C16 |
| C16 | C15 |
| C | O |
| -C | N |

[ impropers ]

| | | | |
|---|---|---|---|
| -C | CA | N | H |
| CA | +N | C | O |
| CZ | NH1 | NE | NH2 |
| C15 | NH2 | C16 | NZ |
| C16 | C15 | NH1 | C20 |

**Table S3.** Atom names, atom types, atom charges, bonds, and impropers for the `LGX` and `AGS` residues of glucosepane crosslink in *GROMACS* `aminoacids.rtp` format and added to the Amber force field alongside the pre-existing entries [1, 2].

##### 4.1.1 Special Bond *(specialbond.dat)*

| resA | atomA | nbondsA | resB | atomB | nbondsB | length | resA | resB |
|---|---|---|---|---|---|---|---|---|
| AGS | NZ | 1 | LGX | CE | 1 | 0.147 | AGS | LGX |

**Table S4.** Special bond definition for the glucosepane crosslink between `LGX-CE` and `AGS-NZ`, reported in *GROMACS* `specbond.dat` format and added to the Amber force field alongside the pre-existing entries [1, 2].

#### 4.2 Pentosidine crosslink

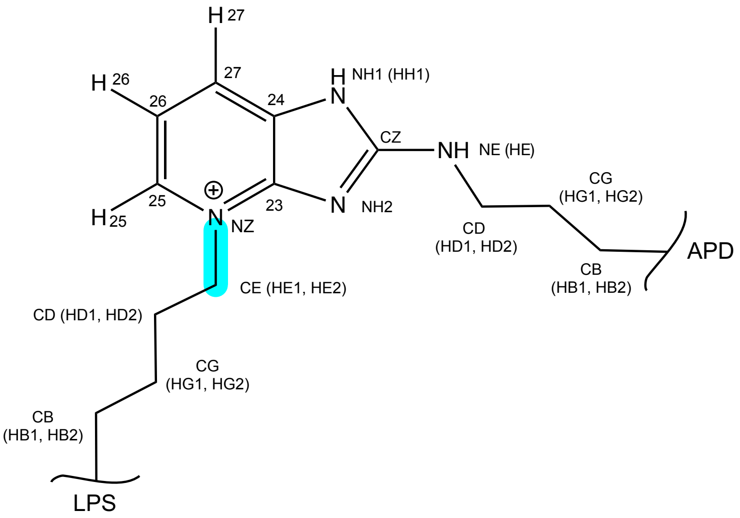


**Figure S3. Pentosidine crosslink.** Atom names and types as defined in the *aminoacids.rtp* file (in *GROMACS* format). The excerpt below shows the entries corresponding to Pentosidine (`LPS`: modified Lys residue & `APD`: modified Arg residue).

**[LPS]**

[ atoms ]

| | | | |
|---|---|---|---|
| N | N | -0.210936 | 1 |
| H | H | 0.194442 | 2 |
| CA | CT | -0.135864 | 3 |
| HA | H1 | 0.106313 | 4 |
| CB | CT | -0.129470 | 5 |
| HB1 | HC | 0.063701 | 6 |
| HB2 | HC | 0.063701 | 7 |
| CG | CT | 0.028893 | 8 |
| HG1 | HC | 0.020343 | 9 |
| HG2 | HC | 0.020343 | 10 |
| CD | CT | 0.016159 | 11 |
| HD1 | HC | 0.029951 | 12 |
| HD2 | HC | 0.029951 | 13 |
| CE | CT | -0.057043 | 14 |
| HE1 | H1 | 0.093913 | 15 |
| HE2 | H1 | 0.093913 | 16 |
| C | C | 0.615994 | 17 |
| O | O | -0.594629 | 18 |

[ bonds ]

| | |
|---|---|
| N | H |
| N | CA |
| CA | HA |
| CA | CB |
| CA | C |
| CB | HB1 |
| CB | HB2 |
| CB | CG |
| CG | HG1 |
| CG | HG2 |
| CG | CD |
| CD | HD1 |
| CD | HD2 |
| CD | CE |
| CE | HE1 |
| CE | HE2 |
| C | O |
| -C | N |

[ impropers ]

| | | | |
|---|---|---|---|
| -C | CA | N | H |
| CA | +N | C | O |

#### [APD]

[ atoms ]

| | | | |
|---|---|---|---|
| N | N | -0.295591 | 1 |
| H | H | 0.255825 | 2 |
| CA | CT | -0.347378 | 3 |
| HA | H1 | 0.186036 | 4 |
| CB | CT | -0.161499 | 5 |
| HB1 | HC | 0.071742 | 6 |
| HB2 | HC | 0.071742 | 7 |
| CG | CT | 0.162532 | 8 |
| HG1 | HC | 0.024357 | 9 |
| HG2 | HC | 0.024357 | 10 |
| CD | CT | -0.075356 | 11 |
| HD1 | H1 | 0.095229 | 12 |
| HD2 | H1 | 0.095229 | 13 |
| NE | N2 | -0.555517 | 14 |
| HE | H | 0.328610 | 15 |
| CZ | CQ | 0.887151 | 16 |
| NH1 | NA | -0.796976 | 17 |
| HH1 | H | 0.481368 | 18 |
| NH2 | NB | -0.699312 | 19 |
| C23 | CM | 0.556462 | 20 |
| C24 | CB | 0.187765 | 21 |
| NZ | N* | -0.077145 | 22 |
| C25 | CM | -0.094801 | 23 |
| H25 | H4 | 0.202281 | 24 |
| C26 | CA | -0.248796 | 25 |
| H26 | HA | 0.214500 | 26 |
| C27 | CA | -0.129208 | 27 |
| H27 | HA | 0.220347 | 28 |
| C | C | 0.737116 | 29 |
| O | O | -0.570742 | 30 |

[ bonds ]

| | |
|---|---|
| N | H |
| N | CA |
| CA | HA |
| CA | CB |
| CA | C |
| CB | HB1 |
| CB | HB2 |
| CB | CG |
| CG | HG1 |
| CG | HG2 |
| CG | CD |
| CD | HD1 |
| CD | HD2 |
| CD | NE |
| NE | HE |
| NE | CZ |
| CZ | NH1 |
| CZ | NH2 |
| NH1 | HH1 |
| NH1 | C24 |
| NH2 | C23 |
| C23 | C24 |
| C23 | NZ |
| NZ | C25 |
| C25 | H25 |
| C25 | C26 |
| C26 | H26 |
| C26 | C27 |
| C27 | H27 |
| C27 | C24 |
| C | O |
| -C | N |

[ impropers ]

| | | | |
|---|---|---|---|
| -C | CA | N | H |
| CA | +N | C | O |
| CZ | NH1 | NE | NH2 |
| C23 | NH2 | C24 | NZ |
| C24 | C23 | NH1 | C27 |

**Table S5.** Atom names, atom types, atom charges, bonds, and impropers for the LPS and APD residues of pentosidine crosslink in *GROMACS* `aminoacids.rtp` format and added to the Amber force field alongside the pre-existing entries [1, 2].

##### 4.2.1 Special Bond *(specialbond.dat)*

| resA | atomA | nbondsA | resB | atomB | nbondsB | length | resA | resB |
|---|---|---|---|---|---|---|---|---|
| LPS | CE | 1 | APD | NZ | 1 | 0.146 | LPS | APD |

**Table S6.** Special bond definition for the pentosidine crosslink between LPS-CE and APD-NZ, reported in *GROMACS* `specbond.dat` format and added to the Amber force field alongside the pre-existing entries [1, 2].

#### 4.3 MOLD crosslink

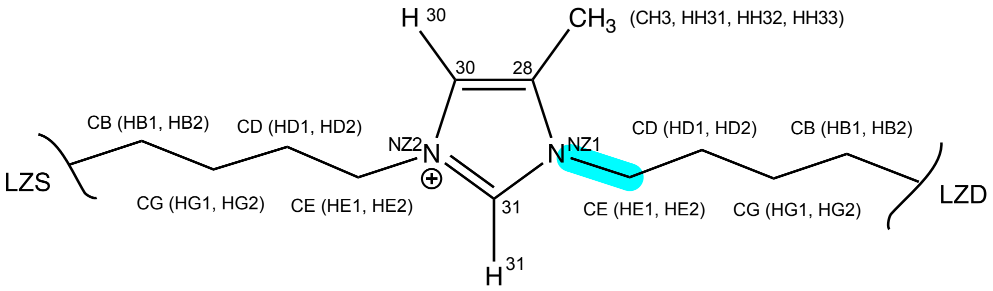


**Figure S4. MOLD crosslink.** Atom names and types as defined in the *aminoacids.rtp* file (in *GROMACS* format). The excerpt below shows the entries corresponding to MOLD (`LZD`: modified Lys & `LZS`: modified Lys).

`[LZD]`

`[ atoms ]`

| | | | |
|---|---|---|---|
| N | N | -0.333459 | 1 |
| H | H | 0.244989 | 2 |
| CA | CT | -0.318983 | 3 |
| HA | H1 | 0.163519 | 4 |
| CB | CT | -0.155066 | 5 |
| HB1 | HC | 0.070095 | 6 |
| HB2 | HC | 0.070095 | 7 |
| CG | CT | 0.083988 | 8 |
| HG1 | HC | 0.059255 | 9 |
| HG2 | HC | 0.059255 | 10 |
| CD | CT | -0.171715 | 11 |
| HD1 | HC | 0.081180 | 12 |
| HD2 | HC | 0.081180 | 13 |
| CE | CT | -0.042829 | 14 |
| HE1 | H1 | 0.100770 | 15 |
| HE2 | H1 | 0.100770 | 16 |
| C | C | 0.702386 | 17 |
| O | O | -0.486187 | 18 |

`[ bonds ]`

| | |
|---|---|
| N | H |
| N | CA |
| CA | HA |
| CA | CB |
| CA | C |
| CB | HB1 |
| CB | HB2 |
| CB | CG |
| CG | HG1 |
| CG | HG2 |
| CG | CD |
| CD | HD1 |
| CD | HD2 |
| CD | CE |
| CE | HE1 |
| CE | HE2 |
| C | O |
| -C | N |

`[ impropers ]`

| | | | |
|---|---|---|---|
| -C | CA | N | H |
| CA | +N | C | O |

#### [LZS]

[ atoms ]

| | | | |
|---|---|---|---|
| N | N | -0.248165 | 1 |
| H | H | 0.192753 | 2 |
| CA | CT | -0.131209 | 3 |
| HA | H1 | 0.152190 | 4 |
| CB | CT | -0.317682 | 5 |
| HB1 | HC | 0.119936 | 6 |
| HB2 | HC | 0.119936 | 7 |
| CG | CT | -0.121262 | 8 |
| HG1 | HC | 0.059129 | 9 |
| HG2 | HC | 0.059129 | 10 |
| CD | CT | 0.179907 | 11 |
| HD1 | HC | 0.006067 | 12 |
| HD2 | HC | 0.006067 | 13 |
| CE | CT | -0.211763 | 14 |
| HE1 | H1 | 0.132006 | 15 |
| HE2 | H1 | 0.132006 | 16 |
| NZ2 | N* | 0.142816 | 17 |
| C | C | 0.638251 | 18 |
| O | O | -0.526574 | 19 |
| C31 | CQ | -0.043104 | 20 |
| H31 | H5 | 0.184965 | 21 |
| NZ1 | N* | -0.066795 | 22 |
| C30 | CC | -0.292230 | 23 |
| H30 | H4 | 0.231915 | 24 |
| C28 | CC | 0.240186 | 25 |
| CH3 | CT | -0.412773 | 26 |
| HH31 | HC | 0.155018 | 27 |
| HH32 | HC | 0.155018 | 28 |
| HH33 | HC | 0.155018 | 29 |

[ bonds ]

| | |
|---|---|
| N | H |
| N | CA |
| CA | HA |
| CA | CB |
| CA | C |
| CB | HB1 |
| CB | HB2 |
| CB | CG |
| CG | HG1 |
| CG | HG2 |
| CG | CD |
| CD | HD1 |
| CD | HD2 |
| CD | CE |
| CE | HE1 |
| CE | HE2 |
| CE | NZ2 |
| C | O |
| -C | N |
| NZ2 | C31 |
| C31 | H31 |
| C31 | NZ1 |
| NZ2 | C30 |
| C30 | H30 |
| C30 | C28 |
| C28 | CH3 |
| CH3 | HH31 |
| CH3 | HH32 |
| CH3 | HH33 |
| C28 | NZ1 |

[ impropers ]

| | | | |
|---|---|---|---|
| -C | CA | N | H |
| CA | +N | C | O |
| NZ2 | C30 | CE | C31 |
| C28 | NZ1 | CH3 | C30 |

**Table S7.** Atom names, atom types, atom charges, bonds, and impropers for the `LZD` and `LZS` residues of MOLD crosslink in *GROMACS* `aminoacids.rtp` format and added to the Amber force field alongside the pre-existing entries [1, 2].

##### 4.3.1 Special Bond *(specialbond.dat)*

| resA | atomA | nbondsA | resB | atomB | nbondsB | length | resA | resB |
|---|---|---|---|---|---|---|---|---|
| LZS | NZ1 | 1 | LZD | CE | 1 | 0.146 | LZS | LZD |

**Table S8.** Special bond definition for the MOLD crosslink between `LZD-CE` and `LZS-NZ1`, reported in *GROMACS* `specbond.dat` format and added to the Amber force field alongside the pre-existing entries [1, 2].

### 5 Newly introduced bonded parameters for AGEs (*ffbonded.itp*)

| **Bonds** | **Atom types** | $\boldsymbol{b_0}$ (nm) | $\boldsymbol{k_b}$ (kJ mol$^{-1}$ nm$^{-2}$) | **Crosslink** |
|---|---|---|---|---|
| NE–CZ | N2–C | 0.13901 | 345010.0 | Glucosepane (AGS) |
| NE–CZ | N2–CQ | 0.13735 | 364180.0 | Pentosidine (APD) |
| CZ–NH1 | CQ–NA | 0.13802 | 356310.0 | Pentosidine (APD) |
| CZ–NH2 | CQ–NB | 0.13694 | 369110.0 | Pentosidine (APD) |
| NH1–C24 | NA–CB | 0.13802 | 356310.0 | Pentosidine (APD) |
| NH2–C23 | NB–CM | 0.13172 | 439650.0 | Pentosidine (APD) |
| C23–C24 | CM–CB | 0.14278 | 351290.0 | Pentosidine (APD) |
| NZ2–C31 | N*–CQ | 0.13710 | 367190.0 | MOLD (LZS) |
| NZ2–C30 | N*–CC | 0.13710 | 367190.0 | MOLD (LZS) |
| C30–H30 | CC–H4 | 0.10830 | 292960.0 | MOLD (LZS) |
| **Angles** | **Atom types** | $\theta_0$ (deg) | $\boldsymbol{k_\theta}$ (kJ mol$^{-1}$ rad$^{-2}$) | **Crosslink** |
| CD–NE–CZ | CT–N2–C | 120.120 | 529.690 | Glucosepane (AGS) |
| HE–NE–CZ | H–N2–C | 115.680 | 405.010 | Glucosepane (AGS) |
| NE–CZ–NH1 | N2–C–N* | 113.640 | 604.170 | Glucosepane (AGS) |
| NH1–CZ–NH2 | N*–C–N* | 120.690 | 606.680 | Glucosepane (AGS) |
| CZ–NH1–C16 | C–N*–C | 116.010 | 571.530 | Glucosepane (AGS) |
| NH2–C15–NZ | N*–CT–N3 | 120.710 | 561.490 | Glucosepane (AGS) |
| NH2–C15–C16 | N*–CT–C | 120.710 | 561.490 | Glucosepane (AGS) |
| H15–C15–C16 | H2–CT–C | 111.690 | 390.790 | Glucosepane (AGS) |
| H15–C15–NZ | H2–CT–N3 | 109.350 | 415.050 | Glucosepane (AGS) |
| NH1–C16–C15 | N*–C–CT | 108.720 | 564.000 | Glucosepane (AGS) |
| CD–NE–CZ | CT–N2–CQ | 119.720 | 533.040 | Pentosidine (APD) |
| HE–NE–CZ | H–N2–CQ | 115.630 | 409.200 | Pentosidine (APD) |
| NE–CZ–NH1 | N2–CQ–NA | 117.280 | 600.820 | Pentosidine (APD) |
| NE–CZ–NH2 | N2–CQ–NB | 117.230 | 603.330 | Pentosidine (APD) |
| NH1–CZ–NH2 | NA–CQ–NB | 121.950 | 590.780 | Pentosidine (APD) |
| CZ–NH1–HH1 | CQ–NA–H | 125.500 | 391.620 | Pentosidine (APD) |
| CZ–NH1–C24 | CQ–NA–CB | 128.010 | 531.370 | Pentosidine (APD) |
| HH1-NH1–C24 | H-NA-CB | 125.500 | 391.620 | Pentosidine (APD) |
| CZ–NH2–C23 | CQ–NB–CM | 105.490 | 600.820 | Pentosidine (APD) |
| NH2–C23–C24 | NB–CM–CB | 112.560 | 599.150 | Pentosidine (APD) |
| NH2–C23–NZ | NB–CM-N* | 112.220 | 626.760 | Pentosidine (APD) |
| C24–C23–NZ | CB–CM–N* | 117.770 | 574.040 | Pentosidine (APD) |
| NH1–C24-C23 | NA–CB–CM | 117.770 | 574.040 | Pentosidine (APD) |
| NH1–C24–C27 | NA-CB-CA | 106.990 | 614.210 | Pentosidine (APD) |
| C23–C24–C27 | CM–CB–CA | 114.190 | 570.700 | Pentosidine (APD) |
| C25–C26–H26 | CM–CA–HA | 121.760 | 405.850 | Pentosidine (APD) |
| CE–NZ2–C31 | CT–N*–CQ | 125.090 | 523.500 | MOLD (LZS,LZD) |
| CE–NZ2–C30 | CT–N*–CC | 125.090 | 523.500 | MOLD (LZS,LZD) |
| C31–NZ2–C30 | CQ–N*–CC | 128.010 | 534.550 | MOLD (LZS) |
| NZ2–C31–H31 | N*–CQ–H5 | 122.100 | 416.390 | MOLD (LZS) |
| NZ2–C31–NZ1 | N*–CQ–N* | 129.110 | 394.890 | MOLD (LZS) |
| NZ2–C30–H30 | N*–CC–H4 | 119.660 | 420.240 | MOLD (LZS) |
| NZ2–C30–C28 | N*–CC–CC | 109.420 | 610.110 | MOLD (LZS,LZD) |
| H30–C30–C28 | H4–CC–CC | 122.780 | 548.100 | MOLD (LZS) |
| NZ1–C28–CH3 | N*–CC–CT | 129.110 | 394.890 | MOLD (LZS) |

| **Proper torsions** | **Atom types** | $\phi_s$ (deg) | $\boldsymbol{k_\phi}$ (kJ mol$^{-1}$) | n | **Crosslink** |
|---|---|---|---|---|---|
| CD–NE–CZ–NH1 | CT–N2–C–N* | 180.00 | 439.320 | 2 | Glucosepane (AGS) |
| HE–NE–CZ–NH2 | H–N2–C–N* | 180.00 | 439.320 | 2 | Glucosepane (AGS) |
| CD–NE–CZ–NH1 | CT–N2–CQ–NA | 180.00 | 4.39320 | 2 | Pentosidine (APD) |
| CD–NE–CZ–NH2 | CT–N2–CQ–NB | 180.00 | 4.39320 | 2 | Pentosidine (APD) |
| HE–NE–CZ–NH1 | H–N2–CQ–NA | 180.00 | 4.39320 | 2 | Pentosidine (APD) |
| HE–NE–CZ–NH2 | H–N2–CQ–NB | 180.00 | 4.39320 | 2 | Pentosidine (APD) |
| NE–CZ–NH1–HH1 | N2–CQ–NA–H | 180.00 | 7.11280 | 2 | Pentosidine (APD) |
| NE–CZ–NH1–C24 | N2–CQ–NA–CB | 180.00 | 7.11280 | 2 | Pentosidine (APD) |
| NH2–CZ–NH1–HH1 | NB–CQ–NA–H | 180.00 | 7.11280 | 2 | Pentosidine (APD) |
| NH2–CZ–NH1–C24 | NB–CQ–NA–CB | 180.00 | 7.11280 | 2 | Pentosidine (APD) |
| NE–CZ–NH2–C23 | N2–CQ–NB–CM | 180.00 | 19.87400 | 2 | Pentosidine (APD) |
| NH1–CZ–NH2–C23 | NA–CQ–NB–CM | 180.00 | 19.87400 | 2 | Pentosidine (APD) |
| NH2–C23–C24–NH1 | NB–CM–CB–NA | 180.00 | 16.73600 | 2 | Pentosidine (APD) |
| CZ–NH1–C24–C23 | CQ–NA–CB–CM | 180.00 | 7.11280 | 2 | Pentosidine (APD) |
| CZ–NH1–C24–C27 | CQ–NA–CB–CA | 180.00 | 7.11280 | 2 | Pentosidine (APD) |
| HH1—NH1–C24–C23 | H–NA–CB–CM | 180.00 | 7.11280 | 2 | Pentosidine (APD) |
| HH1-NH1–C24–C27 | H–NA–CB–CA | 180.00 | 7.11280 | 2 | Pentosidine (APD) |
| CZ–NH2–C23–C24 | CQ–NB–CM–CB | 180.00 | 19.87400 | 2 | Pentosidine (APD) |
| CZ–NH2–C23–NZ | CQ–NB–CM–N* | 180.00 | 19.87400 | 2 | Pentosidine (APD) |
| NH2–C23–C24–C27 | NB–CM–CB–CA | 180.00 | 16.73600 | 2 | Pentosidine (APD) |
| NZ–C23–C24–C27 | N*–CM–CB–CA | 180.00 | 16.73600 | 2 | Pentosidine (APD) |
| NZ–C23–C24–NH1 | N*–CM–CB–NA | 180.00 | 16.73600 | 2 | Pentosidine (APD) |
| CE–NZ2–C31–H31 | CT–N*–CQ–H5 | 180.00 | 7.11280 | 2 | MOLD (LZS,LZD) |
| CE–NZ2–C31–NZ1 | CT–N*–CQ–N* | 180.00 | 7.11280 | 2 | MOLD (LZS,LZD) |
| C30–NZ2–C31–H31 | CC–N*–CQ–H5 | 180.00 | 7.11280 | 2 | MOLD (LZS) |
| C30–NZ2–C31–NZ1 | CC–N*–CQ–N* | 180.00 | 7.11280 | 2 | MOLD (LZS) |
| CE–NZ2–C30–H30 | CT–N*–CC–H4 | 180.00 | 7.11280 | 2 | MOLD (LZS) |
| CE–NZ2–C30–C28 | CT–N*–CC–CC | 180.00 | 7.11280 | 2 | MOLD (LZS,LZD) |
| C31–NZ2–C30–H30 | CQ–N*–CC–H4 | 180.00 | 7.11280 | 2 | MOLD (LZS) |
| C31–NZ2–C30–C28 | CQ–N*–CC–CC | 180.00 | 7.11280 | 2 | MOLD (LZS) |
| NZ2–C31–NZ1–CE | N*–CQ–N*–CT | 180.00 | 7.11280 | 2 | MOLD (LZS,LZD) |
| NZ2–C31–NZ1–C28 | N*–CQ–N*–CC | 180.00 | 7.11280 | 2 | MOLD (LZS) |
| CE–NZ1–C28–CH3 | CT–N*–CC–CT | 180.00 | 7.11280 | 2 | MOLD (LZS,LZD) |
| C31–NZ1–C28–C30 | CQ–N*–CC–CC | 180.00 | 7.11280 | 2 | MOLD (LZS) |
| C31–NZ1–C28–CH3 | CQ–N*–CC–CT | 180.00 | 7.11280 | 2 | MOLD (LZS) |
| NZ2–C30–C28–NZ1 | N*–CC–CC–N* | 180.00 | 7.11280 | 2 | MOLD (LZS) |
| NZ2–C30–C28-CH3 | N*–CC–CC–CT | 180.00 | 16.73600 | 2 | MOLD (LZS) |
| H30–C30–C28–NZ1 | H4–CC–CC–N* | 180.00 | 16.73600 | 2 | MOLD (LZS) |
| H30–C30–C28–CH3 | H4–CC–CC–CT | 180.00 | 16.73600 | 2 | MOLD (LZS) |

| **Improper torsions** | **Atom types** | $\xi_0$ (deg) | $\boldsymbol{k_\xi}$ (kJ mol$^{-1}$ rad$^{-2}$) | n | **Crosslink** |
|---|---|---|---|---|---|
| C16–C15–NH1–C20 | C–CT–N*–CT | 180.00 | 460.240 | 2 | Glucosepane (AGS) |
| CZ–NH1–NE–NH2 | C–N*–N2–N* | 180.00 | 460.240 | 2 | Glucosepane (AGS) |
| C15–NH2–C16–NZ | CT–N*–C–N3 | 180.00 | 460.240 | 2 | Glucosepane (AGS) |
| CZ–NH1–NE–NH2 | CQ–NA–N2–NB | 180.00 | 460.240 | 2 | Pentosidine (APD) |
| C23–NH2–C24–NZ | CM–NB–CB–N* | 180.00 | 460.240 | 2 | Pentosidine (APD) |
| C24–C23–NH1–C27 | CB–CM–NA–CA | 180.00 | 4.60240 | 2 | Pentosidine (APD) |
| NZ2–C30–CE–C31 | N*–CC–CT–CQ | 180.00 | 460.240 | 2 | MOLD (LZS,LZD) |
| C28–CH3–NZ1–C30 | CC–CT–N*–CC | 180.00 | 460.240 | 2 | MOLD (LZS,LZD) |

**Table S9.** Bonded parameters for the AGE crosslinks, written in *GROMACS* format, were added to the *ffbonded.itp* file alongside those reported in [1, 2].

#### 6 Benchmarking AGEs Crosslink Force Field Parameters

AGE structures obtained after *ab initio* geometry relaxation were solvated in a 6 nm cubic box of TIP3P water [3] and neutralized with NaCl. Following energy minimization, systems were equilibrated for 10 ns in the NVT ensemble with heavy-atom restraints on the crosslink, then for 15 ns in the NPT ensemble without restraints. Temperature (300 K) and pressure (1 bar) were controlled with the velocity-rescale thermostat [4] and the Parrinello–Rahman barostat [5], and electrostatics were treated with PME [6]. For each crosslink, five independent 100-ns replicas were run with each parameter set (1 µs per crosslink; 3 µs total) using GROMACS 2024.4 [7]. Table S3 reports the ring-angle values for each AGE in the QM-optimized reference structure together with the corresponding replica-averaged values from MD simulations carried out with the two parameter sets, as well as the associated uncertainty, reported as the standard deviation (SD) of the angle distribution across trajectory frames. Angle definitions follow the atom nomenclature in Figures S2–S4. Angles involving bonds between ring atoms and side-chain atoms and/or ring substituents (C–OH and C–$CH_3$ substituents in glucosepane and MOLD, respectively) are highlighted in yellow. Figures S5 presents RMSD, RMSF, and end-to-end distance (E2E) structural metrics evaluated relative to the QM structure.

| **Glucosepane** | | | | | | | |
|---|---|---|---|---|---|---|---|
| *Angle (deg)* | *QM* | *Antechamber* | *Grappa* | *Angle (deg)* | *QM* | *Antechamber* | *Grappa* |
| NH1−CZ−NH2 | 117.25 | 115.20 ± 2.7 | 116.28 ± 2.05 | C19−C20−C16 | 108.76 | 114.43 ± 3.15 | 111.21 ± 3.62 |
| NH2−CZ−NE | 125.10 | 111.28 ± 4.70 | 121.38 ± 2.72 | C18−C19−C20 | 115.89 | 114.17 ± 3.60 | 114.16 ± 3.23 |
| NH1−CZ−NE | 117.63 | 116.79 ± 4.33 | 122.03 ± 2.64 | C18−C19−O19 | 109.11 | 109.42 ± 3.69 | 109.06 ± 3.14 |
| C16−NH1−CZ | 103.89 | 108.42 ± 2.63 | 104.98 ± 2.08 | C20−C19−O19 | 107.85 | 107.12 ± 3.68 | 109.15 ± 3.10 |
| C15−NH2−CZ | 103.88 | 103.11 ± 2.52 | 103.82 ± 2.07 | C17−C18−C19 | 117.03 | 114.88 ± 3.76 | 114.32 ± 3.31 |
| NH2−C15−C16 | 103.67 | 106.99 ± 2.42 | 104.02 ± 1.97 | C17−C18−O18 | 111.47 | 105.06 ± 3.77 | 108.49 ± 3.21 |
| NZ−C15−NH2 | 117.84 | 118.78 ± 3.24 | 115.96 ± 3.40 | C19−C18−O18 | 107.71 | 110.13 ± 3.66 | 108.82 ± 3.15 |
| NZ−C15−C16 | 112.93 | 111.75 ± 2.92 | 112.80 ± 3.31 | NZ−C17−C18 | 111.40 | 112.63 ± 3.22 | 115.12 ± 3.90 |
| C15−C16−NH1 | 110.82 | 104.27 ± 2.52 | 110.40 ± 2.03 | C17−NZ−CE | 118.19 | 112.10 ± 3.84 | 117.29 ± 3.62 |
| C20−C16−C15 | 124.0 | 124.61 ± 3.19 | 123.24 ± 3.11 | C15−NZ−CE | 114.47 | 116.10 ± 3.96 | 114.76 ± 3.58 |
| **Pentosidine** | | | | | | | |
| *Angle (deg)* | *QM* | *Antechamber* | *Grappa* | *Angle (deg)* | *QM* | *Antechamber* | *Grappa* |
| NH1−CZ−NH2 | 112.68 | 116.74 ± 2.28 | 112.61 ± 1.99 | C23−C24−C27 | 121.63 | 122.77 ± 2.27 | 122.55 ± 2.42 |
| NE−CZ−NH1 | 122.84 | 121.94 ± 3.17 | 120.55 ± 2.83 | C24−NH1−CZ | 107.32 | 101.52 ± 2.20 | 106.53 ± 1.97 |
| NH2−CZ−NE | 124.46 | 119.97 ± 3.26 | 126.49 ± 2.69 | C24−C27−C26 | 116.63 | 115.26 ± 2.28 | 114.74 ± 2.45 |
| CZ−NH2−C23 | 104.99 | 100.79 ± 2.14 | 104.74 ± 1.99 | C27−C26−C25 | 119.79 | 120.86 ± 2.27 | 122.02 ± 2.47 |
| NH2−C23−C24 | 112.15 | 110.59 ± 2.11 | 112.54 ± 1.99 | C26−C25−NZ | 122.49 | 120.73 ± 2.22 | 121.31 ± 2.37 |
| C25−NZ−C23 | 118.86 | 117.14 ± 2.18 | 118.73 ± 2.37 | C25−NZ−CE | 120.88 | 120.26 ± 2.62 | 121.43 ± 2.82 |
| C24−C23−NZ | 120.56 | 122.03 ± 2.18 | 119.70 ± 2.37 | C23−NZ−CE | 120.25 | 121.96 ± 2.59 | 119.27 ± 2.84 |
| C23−C24−NH1 | 102.82 | 109.74 ± 2.17 | 103.12 ± 1.957 | | | | |
| **MOLD** | | | | | | | |
| *Angle (deg)* | *QM* | *Antechamber* | *Grappa* | *Angle (deg)* | *QM* | *Antechamber* | *Grappa* |
| NZ2−C31−NZ1 | 109.35 | 106.77 ± 2.34 | 108.17 ± 2.01 | C30−C28−CH3 | 129.43 | 121.48 ± 2.85 | 129.28 ± 3.18 |
| C31−NZ1−C28 | 108.5 | 109.79 ± 2.25 | 109.29 ± 2.01 | C28−C30−NZ2 | 108.62 | 104.51 ± 2.11 | 107.16 ± 2.03 |
| C31−NZ1−CE | 123.18 | 124.08 ± 2.88 | 123.26 ± 3.17 | C30−NZ2−C31 | 107.65 | 111.40 ± 2.27 | 108.89 ± 2.05 |
| CE−NZ1−C28 | 128.29 | 125.43 ± 2.84 | 126.79 ± 3.12 | C30−NZ2−CE | 124.68 | 123.74 ± 2.96 | 125.39 ± 3.28 |
| NZ1−C28−C30 | 105.86 | 106.81 ± 2.11 | 106.13 ± 2.01 | C31−NZ2−CE | 127.65 | 124.21 ± 2.95 | 125.22 ± 3.27 |
| NZ1−C28−CH3 | 124.70 | 130.65 ± 2.81 | 124.22 ± 3.07 | | | | |

**Table S10. Ring-angle geometry of AGE crosslinks.** QM values are taken from the DFT-optimized reference structures; MD values (Antechamber and Grappa) are reported as mean ± standard deviation over trajectory frames. Angle definitions follow Figures S2–S4. Angles highlighted in yellow involve bonds between ring atoms and side-chain atoms and/or ring substituents and are shown for completeness but were not included in the ring-core benchmarking discussed in the main text.

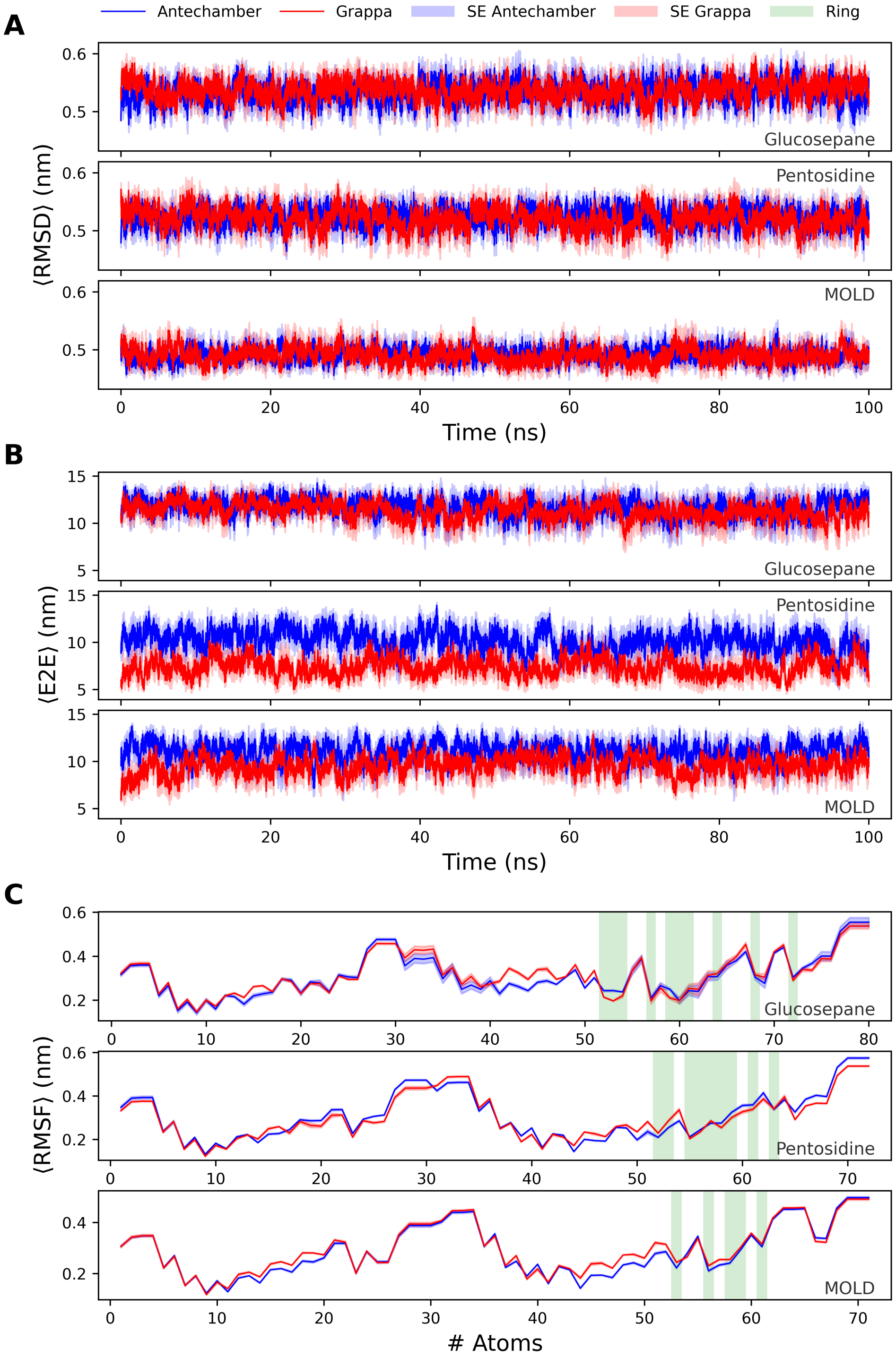


**Figure S5. Structural metrics of AGE crosslinks from MD simulations.** Blue and red curves correspond to the Antechamber and Grappa parameter sets, respectively; shaded regions indicate the standard error of the mean (SEM) across replicas. **(A)** ⟨RMSD⟩ of the crosslink (all atoms) relative to the QM reference structure as a function of time. **(B)** ⟨E2E⟩, defined as the distance between the two crosslink C$\alpha$ atoms, as a function of time. **(C)** ⟨RMSF⟩ (all atoms) relative to the QM reference structure; green bands highlight ring atoms. Time-averaged values are reported in Table S11.

| Structural metrics | Crosslink type | Antechamber | Grappa |
|---|---|---|---|
| | Glucosepane | $0.533 \pm 0.015$ | $0.537 \pm 0.015$ |
| $\langle$RMSD$\rangle$ (nm) | Pentosidine | $0.524 \pm 0.013$ | $0.521 \pm 0.017$ |
| | MOLD | $0.491 \pm 0.010$ | $0.490 \pm 0.012$ |
| | Glucosepane | $0.124 \pm 0.001$ | $0.126 \pm 0.001$ |
| $\langle$RMSD$\rangle_{\text{ring}}$ (nm) | Pentosidine | $0.139 \pm 0.001$ | $0.135 \pm 0.001$ |
| | MOLD | $0.129 \pm 0.001$ | $0.129 \pm 0.001$ |
| | Glucosepane | $11.700 \pm 0.847$ | $11.321 \pm 0.853$ |
| $\langle$E2E$\rangle$ (nm) | Pentosidine | $10.154 \pm 0.919$ | $7.356 \pm 0.824$ |
| | MOLD | $11.066 \pm 0.912$ | $9.523 \pm 0.861$ |
| | Glucosepane | $0.310 \pm 0.011$ | $0.317 \pm 0.008$ |
| $\langle$RMSF$\rangle$ (nm) | Pentosidine | $0.308 \pm 0.004$ | $0.302 \pm 0.004$ |
| | MOLD | $0.287 \pm 0.003$ | $0.297 \pm 0.003$ |
| | Glucosepane | $0.254 \pm 0.009$ | $0.242 \pm 0.009$ |
| $\langle$RMSF$\rangle_{\text{ring}}$ (nm) | Pentosidine | $0.277 \pm 0.003$ | $0.275 \pm 0.005$ |
| | MOLD | $0.297 \pm 0.003$ | $0.269 \pm 0.002$ |

**Table S11. Time-averaged structural metrics for AGE crosslinks.** Values are reported as mean $\pm$ SEM across replicas. Blue/red: Antechamber/Grappa. $\langle$RMSD$\rangle$ (Figure S5A) is computed relative to the QM reference structure; $\langle$RMSD$\rangle_{\text{ring}}$ and $\langle$RMSF$\rangle_{\text{ring}}$ are computed using ring atoms only; $\langle$E2E$\rangle$ (Figure S5B) is the C$\alpha$–C$\alpha$ distance between the two crosslinking residues; $\langle$RMSF$\rangle$ (Figure S5C) quantifies per-atom flexibility around the mean structure.

### 7 Collagen microfibril all-atom simulations

Geometric observables were computed from *mean positions* (centers of geometry) of predefined atom groups. The end-to-end distance was calculated for each frame as the Euclidean distance between the mean positions of the ACE (N-terminus) and NME (C-terminus) cap residues, and is reported as a proxy for the molecular extension (D-band length).

To partition the extension into overlap and gap contributions, we identified the crosslink sets located at the N- and C-terminal sites and used their positions for per-frame clustering. For the PYD and PYD + glucosepane systems, enzymatic crosslinks were detected from the bond network as connected components restricted to the trivalent marker residues (`LYX`, `LY2`, `LY3`), retaining only valid trios containing one residue of each type. For the AGE-terminated systems, crosslinks were instead represented as marker *pairs* and clustered accordingly: `AGS`/`LGX` for glucosepane, `LPS`/`APD` for pentosidine, and `LZS`/`LZD` for MOLD. In all cases, a crosslink position in a given frame was defined as the mean position of all atoms belonging to the corresponding marker set (trio for PYD; pair for AGEs).

For each frame, marker-set mean positions were projected onto the instantaneous molecular axis defined by the ACE→NME direction. The resulting one-dimensional coordinates were segmented using $k = 2$ K-means clustering, yielding two clusters corresponding to crosslinks located closer to the N-terminus and C-terminus, respectively. The *overlap* length was defined as the axial separation between the two cluster centroids, and the *gap* length as the residual extension (gap = end-to-end − overlap). This two-cluster segmentation is appropriate for the present reduced system length, where only two terminal crosslink groups are resolved along the fibril axis.

Uncertainties for overlap and gap were estimated from the within-cluster spread of projected crosslink coordinates by computing standard errors of the cluster means and propagating them to the overlap (difference of cluster centroids) and gap (end-to-end minus overlap). For the end-to-end distance, we combined (i) the propagated frame-wise contribution arising from overlap and gap uncertainties with (ii) a trajectory-scale bootstrap estimate obtained using a 10-frame sliding window and 1000 resamples per window; the two contributions were combined by a root-mean-square procedure. The overlap/gap strain ratio was computed relative to the first analyzed frame as $(Overlap(t) - Overlap(t_0))/(Gap(t) - Gap(t_0))$, with uncertainty propagated from the corresponding overlap and gap errors. Three independent replicas with distinct initial velocities were analyzed. Replica-averaged time series were obtained by arithmetic averaging, and uncertainties on the averaged curves were computed by combining per-replica errors in quadrature and scaling by the number of replicas.

### 8 Extending the ColBuilder framework with new crosslinks

This section provides a practical guide for users who wish to add crosslink chemistries that are not yet included in the ColBuilder database. We outline the minimal set of edits required to (i) register a new crosslink type for model building and geometry refinement and (ii) enable all-atom topology export by providing compatible Amber99 force-field parameters.

**Crosslink representation and database entry.** In ColBuilder, each crosslink is encoded using custom *marker residues* that represent the chemically modified amino acids participating in bond formation. Divalent crosslinks are represented by two markers, whereas trivalent crosslinks are represented by three. How the underlying chemistry is partitioned across markers is a modeling choice (e.g., assigning a ring system to one marker), but the pipeline requires that marker identities and their forming atoms are defined consistently.

New crosslinks are registered by adding one row per crosslink definition to `colbuilder/src/colbuilder/data/sequence/crosslinks.csv` (see Table S1 for an example). In practice, entries are selected by matching (i) the `term` flag (`N` or `C`), (ii) the `type` label (the crosslink name selected in the configuration), and (iii) the `combination` string (a user-defined key matched exactly). Each row specifies the marker residues and their forming atoms via `R1/A1/P1`, `R2/A2/P2` (and `R3/A31/P3` for trivalent links), where positions `P` are given as `resnum.chain` (e.g., `1004.B`). The `#shift` field is retained as an annotation to record the unit-cell translation required to optimize the crosslink.

**MODELLER residue registration.** Each marker must be available to MODELLER as a residue type. Concretely, markers are mapped to single-character residue identifiers in `colbuilder/src/colbuilder/data/sequence/modeller/restyp_mod.lib`, and their heavy-atom topology is added to `colbuilder/src/colbuilder/data/sequence/modeller/top_heav_mod.lib`. To streamline the latter step, we provide a helper script in `https://github.com/gggguido/AGEs-ColBuilder.git` that generates a copy/paste-ready topology block for `top_heav_mod.lib` starting from a marker geometry specified as a SMILES string. The script (`create_ic_crosslink.py`) builds a 3D conformer with RDKit, assigns systematic atom naming, and writes an internal-coordinate (IC) specification together with the required bonded connectivity, reducing manual editing when defining additional markers.

**Geometry pipeline integration.** To ensure that the geometry and refinement routines recognize the new markers, the marker lists and crosslink parsing logic must be extended in `colbuilder/src/colbuilder/core/geometry/crosslink.py` and `colbuilder/src/colbuilder/core/geometry/connect.py`. In addition, the pairing/trio rules and replacement mappings that support crosslink mixing, correction of unpaired markers, and user-controlled marker replacement must be updated in `colbuilder/src/colbuilder/core/geometry/geometry_replacer.py`, `colbuilder/src/colbuilder/core/geometry/unpaired_crosslinks.py`, and `colbuilder/src/colbuilder/core/geometry/crosslink_mixer.py`. Collectively, these files define which marker residues are considered compatible within a crosslink and how markers are reverted to their corresponding standard residues (e.g., Lys/Arg) when required by specific workflow operations.

**All-atom parameterization (required for topology export).** If the new markers are intended for all-atom molecular dynamics, they must be parameterized and made available to the topology builder. We followed a standard AMBER/GAFF workflow, using AmberTools [8], based on quantum-mechanical geometry optimization and electrostatic-potential (ESP) fitting. For each marker, we ran a Gaussian calculation at the B3LYP/6-31G* level to optimize the geometry and compute a Merz–Kollman ESP (via `Pop=MK` and the associated `iop` settings), yielding the Gaussian output and an ESP grid (`.gesp`). The ESP grid was converted to a `.esp` file using `espgen` and used for a two-stage RESP fit: starting from the Gaussian output, `antechamber` generated an initial `.ac` file, respgen prepared respin1/respin2 (and qin), and RESP stage 1 and stage 2 fits produced the final restrained charge set. In the stage 2 fit, charges on the acetyl (ACE) and N-methyl (NME) caps were constrained to reproduce the corresponding AMBER values by freezing the cap-atom charges through the qin/constraint specification used by respgen. The resulting RESP charges were assigned back to the structure with antechamber; GAFF atom types were then applied, and missing bonded parameters were identified with parmchk to generate the .frcmod file. Finally, .mol2 (RESP charges) and .prepi/.frcmod files were produced for LEaP, enabling generation of AMBER topology/coordinate files (.prmtop/.inpcrd) for subsequent conversion via acpype and use in *GROMACS* simulations. A full example is reported in https://github.com/gggguido/AGEs-ColBuilder.git.

**All-atom topology generation (Amber99SB*-ildnp).** For ColBuilder to export all-atom topologies, the topology layer must recognize each marker residue and its bonding atom. This mapping is defined in colbuilder/src/colbuilder/core/topology/amber.py. The force-field data files then provide atom types, partial charges, hydrogen-generation rules, residue classification, and special-bond definitions. For Amber99sb*-ildnp, these edits are made in colbuilder/src/colbuilder/data/topology/amber99sb-star-ildnp.ff/, specifically in aminoacids.rtp (residue definitions), specbond.dat (special crosslink bonds), residuetypes.dat (residue classification), and aminoacids.hdb (hydrogen construction rules). If Amber99 output is also required, the corresponding residue definition must additionally be added in colbuilder/src/colbuilder/data/topology/amber99/ (file: aminoacids.rtp).
